# Comparable anti-ageing efficacies of a multi-ingredient nutraceutical and a senolytic intervention in old mice

**DOI:** 10.1101/2024.10.11.617853

**Authors:** Charlotte Brookes, Edward Fielder, Evon Low, Diogo Barardo, Thomas von Zglinicki, Satomi Miwa

## Abstract

Single-ingredient dietary supplements have demonstrated some potential to extend lifespan and improve healthspan; however, the efficacy of defined multi-ingredient nutraceuticals remains underexplored. Senolytic interventions have been successful in reducing multi-morbidity, frailty, and cognitive decline in animal models and are seen as promising anti-ageing interventions in mammals. We compared the effects of a 12-ingredient nutraceutical on mice lifespan and healthspan markers with that of a high efficacy senolytic intervention consisting of a low dose of Navitoclax combined with the specific mitochondrial uncoupler BAM15. Both interventions were started at old age (20 months). The supplement was given daily until the end of the experiment (30 months of age), but the senolytic intervention consisted of two short (5 days each) rounds of treatment at 20 and 23 months of age. Despite late onset, both interventions increased median lifespan similarly by around 20% over controls. The senolytic intervention significantly reduced frailty progression and improved cognitive function after the second round of treatment, but without subsequent treatment, these effects appeared to wane at later ages. The multi-ingredient supplement tended to reduce frailty progression steadily with time, albeit not significant, and maintained cognitive function. Mechanistically, *in vitro*, there was no evidence of senolytic activity of the multi-ingredient nutraceutical as a whole nor of its individual ingredients. Continuous multi-ingredient dietary supplementation shows promise in achieving comparable anti-ageing efficacy as a senolytic intervention.

## Introduction

For an ever increasing range of nutraceuticals (dietary supplements), beneficial effects on lifespan and (markers of) healthspan have been demonstrated in multiple organisms^1-8^, some of which have even been tested in clinical trials; including fisetin^9^, pterostilbene^10^ or alpha-ketoglutarate^11^. Given that nutraceuticals typically aim to target multiple overlapping mechanisms of ageing, which are often interlinked^12^, it’s plausible to assume additive or even synergistic health- and lifespan effects of nutraceuticals. While food-derived mixtures of polyphenols led to improvements in lipid and inflammation status following short-term, high-dose interventions in humans^13^, the effects of combining multiple low-dose nutraceuticals has seldom been evaluated. The NOVOSLabs company has recently developed a supplement combining low doses of 12 nutraceutical ingredients with individually demonstrated anti-ageing efficacy, namely pterostilbene^7^, glucosamine sulfate^2^, fisetin^6^, glycine^1^, lithium aspartate^4^, calcium alpha-ketoglutarate^8^, magnesium malate^14^, vitamin C (ascorbic acid)^15^, L-theanine^16^, hyaluronic acid^17^, Rhodiola rosea root extract^18^, and ginger root extract^19^. This supplement has recently shown potential as an antioxidant and for reducing DNA damage *in vitro*^20^, however, its effects on mammalian lifespan or healthspan are unknown.

In recent years, drugs that specifically ablate senescent cells (senolytics) have become the promising anti-ageing interventions in mammals^21^. First generation senolytics including Navitoclax or the combination of Dasatinib and Quercetin were extremely successful in promoting healthy longevity in a wide range of mouse models for age-associated diseases and disabilities^22^. We recently identified mitochondrial membrane potential as a specific vulnerability of senescent cells than can be targeted by mitochondrial uncouplers, and showed that combination with a mitochondrial uncoupler (e.g. BAM15) increased the senolytic activity of BH3 mimetics, such as Navitoclax, by about two orders of magnitude and rescued premature ageing phenotypes induced by sublethal whole-body irradiation in mice^23^. This approach has not been examined in naturally aged mice.

Here, we compare the efficacies of the NOVOS multi-ingredient nutraceutical and the senolytic BAM15/Navitoclax treatment^23^ to improve lifespan and healthspan indicators in old mice, aged 20 months at start of the interventions. A continuous nutraceutical intervention increased lifespan of animals as effectively as two short bouts of senolytic at 20 and 23 months of age. Ongoing nutraceutical intervention caused continuous improvements of the frailty index and better maintained short-term memory as assessed by spontaneous alternation in a Y maze test over controls. In contrast, improvements in both frailty index and short-term memory caused by the senolytic were lost at 4 months after treatment.

## Material and Methods

### Animals

38 C57Bl/6J male mice were bought past weaning from Charles River. The mice were housed in same-sex cages in groups of 3 (56 × 38 × 18 cm, North Kent Plastics, Kent, UK) and were identified by ear notches. They were provided with sawdust, paper bedding and environmental enrichment (cardboard tunnels). Mice were housed at 20 ± 2°C under a 12h light/12h dark photoperiod. The work was licensed by the UK Home Office (PBDAFDFB0) and complied with the guiding principles for the care and use of laboratory animals. Mice were monitored daily and were humanly killed if they displayed severe distress and/or persistent weight loss according to the approved study plan (Supplementary Table S1).

At 18-months old, the mice were ranked on frailty score (low to high), cognitive function (high to low) and body weight (high to low). Ranks were averaged for each mouse, and the animals were assigned to each experimental group; Control (C), Dietary Supplement (DS), and Senolytic (SEN) according to their overall ranked scores, in order to reduce variability between the groups. Treatment began at 20 months old, with frailty and cognition assessed every other month, at least 4-weeks after the final day of drug treatment, to allow the mice to recover following oral gavage and injections.

### Interventions

The multi-ingredient dietary supplement was produced by the company NOVOSLabs. Ingredients and weight per recommended serving for a 70kg human are listed in supplementary Table S2. The supplement was administered to the mice in soaked food, prepared as follows. Food pellets (CRM-P formulation rodent diet, SDS diets) were ground into a powder, mixed with the supplement, and then made as ‘soaked food’ using sterile water at a 1:2 ratio to form a ‘porridge-like’ consistency. The amount of supplement added to the soaked food was determined based on a small dose-finding study (see below). Soaked food (with and without supplement) was given fresh every day of the study, starting when the mice were 20-months old.

Senolytic treatment was administered as described^23^. Briefly, mice assigned to the SEN group were orally gavaged with 0.5mg/kg BW Navitoclax in 10% Polyethylene Glycol (PEG) 400 for 5 consecutive days. They were simultaneously injected with 2.5mg/kg BW BAM15 in 40% PEG400. In parallel, all other mice were gavaged and injected with vehicle only.

Navitoclax (A3007-APE-100mg) was purchased from Stratech, BAM15 (Cat. 5737) was purchased from Biotechne, and PEG400 (8074851000) was purchased from Merck. Two months after the first course of treatment, the second and final course was administered again for five days. During the first course, BAM15 was intraperitoneally injected, however, due to high rates of infection, the injection route was altered to subcutaneous for the second course.

### Mouse Assessments

Frailty was assessed using a 30-parameter scoring index, adapted from Whitehead et al., 2014^24^, during the light photoperiod. To reduce the stress on the mice, we did not test the menace reflex, instead forelimb grip strength was measured using BIO-GS3 BIOSEP. Forelimb strength was recorded 3-times per mouse, the average of which was compared to reference strength values from sex-matched adult mice.

Cognitive function was assessed using a Spontaneous Alternation protocol with a Y-Maze^24^, during the dark photoperiod. The Y-maze was made of dark grey plastic, each arm 40cm long, 5cm wide, and 10cm high. Mice were placed in arm 1 and were allowed to freely explore the maze for 8 minutes in near-dark light conditions. During the 8 minutes, the order of maze arm entry was manually recorded. A ‘spontaneous alternation’ is defined by a mouse entering a different arm of the maze in 3 consecutive entries. Spontaneous alternation frequency is then calculated as the number of spontaneous alternations, divided by the total number of arm entries minus 2. Mice that appear anxious can be observed to prefer the home arm and not explore the whole maze rendering spontaneous alternation measurement uninformative. Mice that spent the majority of their time in the home arm (entered the home arm in more than half of all arm entries) were excluded (2 mice total).

### Cell viability assay

Cell viability of senescent vs non-senescent cells was measured as described^23^. Briefly, cells were senesced using 20Gy X-Ray irradiation *in situ* on 96 well plates, and cultured for 10 days prior to drug treatment. Proliferating cells were plated 24 hours prior to drug exposure. Serial dilutions of compounds were performed to give dose-response curves. Solubility of compounds varied, and DMSO/Ethanol/H2O were used depending on the compound, with the solvent percentage equal across all dilutions (supplementary Table S3). Cells were treated with compound of interest for 72 hours, and stained with crystal violet (0.2% crystal violet/1% ethanol) for 10 minutes, washed with water and dried, solubilised with 1% SDS and absorbance read at 590nm on a plate reader (FLUOstar Omega, BMG).

### Senolysis and cell size assay

Cell numbers and average sizes of human diploid fibroblasts (in senescent/non-senescent co-culture) following treatment with the multi-ingredient supplement were measured as described^23^. Briefly, cells were pre-labelled with whole-cell pSLIEW or mCherry by stable transduction prior to experiments. pSLIEW labelled cells were senesced *in situ* using 20Gy X-Ray irradiation 10 days prior to drug treatment. Proliferating mCherry labelled cells were added into co-culture 1 day prior to drug treatment. Co-cultures were treated with complete commercial NOVOS multi-ingredient supplement for 72 hours. Stock solutions of multi-ingredient supplement were prepared in water and ethanol due to differing solubility of constituent compounds. Cells are imaged prior to, and post-treatment (Leica DMi-8 with automated stage). The area covered by senescent and non-senescent cells was measured by thresholding in ImageJ. This was divided by the respective number of cells in the field, to give average cell size per field. A decrease in relative number of senescent vs non-senescent cells indicated senolytic activity of the tested compound, while a decrease in senescent cell size is indicative of a potential senostatic effect. Rapamycin was used as a positive control (Supplementary Fig. S1).

### Statistics

All statistics and graphics were performed using GraphPad Prism 10.3. One-way and two-way analysis of variance, and nonlinear regression analysis tests were performed using GraphPad Prism 10.3. Survival was assessed using Kaplan-Meyer curves and log-rank analysis.

## Results

### 1. Dietary Supplement Dose Optimisation

Food intake in 20 young male C57Bl/6J mice was measured for 8 days before and 7 days after switching from dry to soaked food. Food intake increased to an average of 5.3g/day from 2.6g/day (Supplementary Fig. S2a), and this value was used to calculate the amount of the multi-component supplement added into the soaked food to achieve 1x, 5x, 10x, and 25x the recommended daily human dose per kg body weight (BW). Four mice per group were fed with these doses (the control group received soaked food without added supplement), and body weight (BW) changes were monitored daily over 14 days (Supplementary Fig. S2b). There were significant effects for time and animal, but not for dose (p=0.312). After 14 days, there were no differences between groups in body weight, grip strength, or body temperature (not shown). Necropsy revealed increased fat depots in all groups but no signs of pathology and no differences in liver, spleen, or kidney weights between groups. Concluding that the supplement was well tolerated by the mice up to 25x the recommended human dose. A dose of 12.3x was chosen because this had been suggested as the appropriate conversion factor from humans to mice^25^. The composition of the dietary supplement at 12.3x dose for consumption of a 40g mouse can be found in Supplementary Table S4.

## 2. A multi-ingredient nutraceutical intervention improved survival as much as two rounds of senolytic

At 18 months of age, male mice were assessed and ranked based on a combined score for frailty, neuromuscular performance, cognitive performance, and body weight at this pre-treatment baseline. Given that there is substantial variation between mice at this age, the mice were distributed to treatments based on this ranking to ensure minimal pre-existing variation between each group. These were Control (C), Dietary Supplement (DS) and Senolytic (SEN) (experimental overview - Fig. 1a.). All animals received soaked food from 20 months of age. Food was supplemented with 0.87% w/w NOVOS nutraceuticals (e.g. 4.338g in 500g) for the DS group: starting at 20 months of age and continuing over the whole experiment. Mice in the SEN group received 0.5mg/kg BW Navitoclax (by gavage) and 2.5mg/kg BW BAM15 (per injection) for 5 consecutive days at months 20 and 23. On the same days, DS and Control mice were also gavaged and injected with control solutes. Mice were humanely killed when showing signs of severe distress (Supplementary Table S1). Deaths occurring before beginning of treatment (2 deaths in the SEN group) were censored, and follow-up was terminated at an age of 896 days. A Kaplan-Meier analysis showed significant improvements in survival for both interventions (Fig. 1b). Median survival [±95% CI] increased from 703 [637,720] days in controls to 830 [645,868] days in the DS group and to 834 [637,889] days in the SEN group. Increased longevity was also reflected in longitudinal body weight data (Fig. 1c): During the period before the first assessment at 21 months, almost all mice displayed increasing BW, evidently due to increased food intake after the switch to soaked food. However, after 21 months of age BW stagnated or decreased in control mice while it still increased in most of the treated animals until 24 or even 27 months of age. Retention of body mass, and especially fat mass, at higher age has been shown to be a predictor of survival in mice^26-28^.

**Fig. 1.**
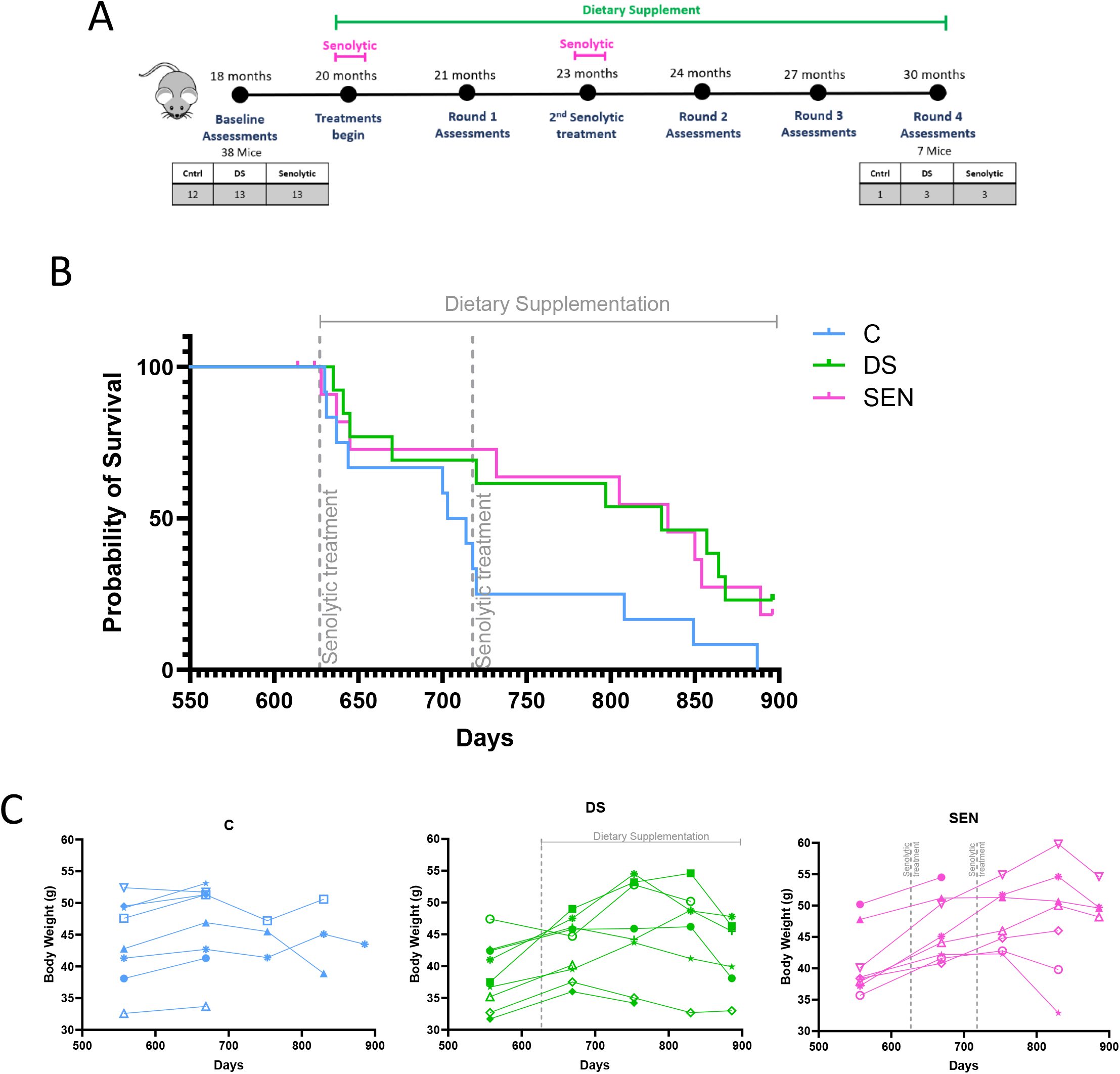
Effects of a multi-ingredient dietary supplement (DS) and a senolytic (SEN) on survival in male mice. **A)** Design of the experiment. **B)** Kaplan-Meier survival curves. Log-rank tests for the difference to controls resulted in p=0.045 (DS) and p=0.043 (SEN), respectively. **C)** Body weights of individual mice. Left (blue): controls (C), middle (Green): dietary supplement (DS), right (pink): Senolytic (SEN) group).

### 3. Improvements in frailty and cognition depend on length of intervention

Frailty indices measured in the present study Control mice fitted closely with those measured in untreated ageing male C57Bl/6J previously^24^ (Fig. 2a). Both DS and SEN interventions reduced the progression of frailty in treated mice, however with different patterns: the continuously provided supplement diminished frailty progression more with ongoing time of intervention, almost reaching statistical significance at the 30 months assessment compared with controls (Fig. 2b). In contrast, the efficacy of the senolytic intervention was greatest shortly after the second round of intervention (assessment round 2) and waned afterwards (Fig. 2c). Two rounds of senolytic treatment improved short-term memory assessed as spontaneous alternation in a Y maze (assessment round 2) (Fig. 2d), whereas continuous dietary supplementation sustained short-term memory, meanwhile the control short-term memory declined as time progressed (Fig 2d). To note, assigning the groups using combined ranks of frailty, BW and spontaneous alternation score resulted in a lower average memory score for the DS group at baseline, however this was improved to control levels, but not beyond, by the treatment (Supplementary Fig. S3). There were insufficient number of animals in control groups to compare the groups at round 3 and 4.

**Fig. 2.**
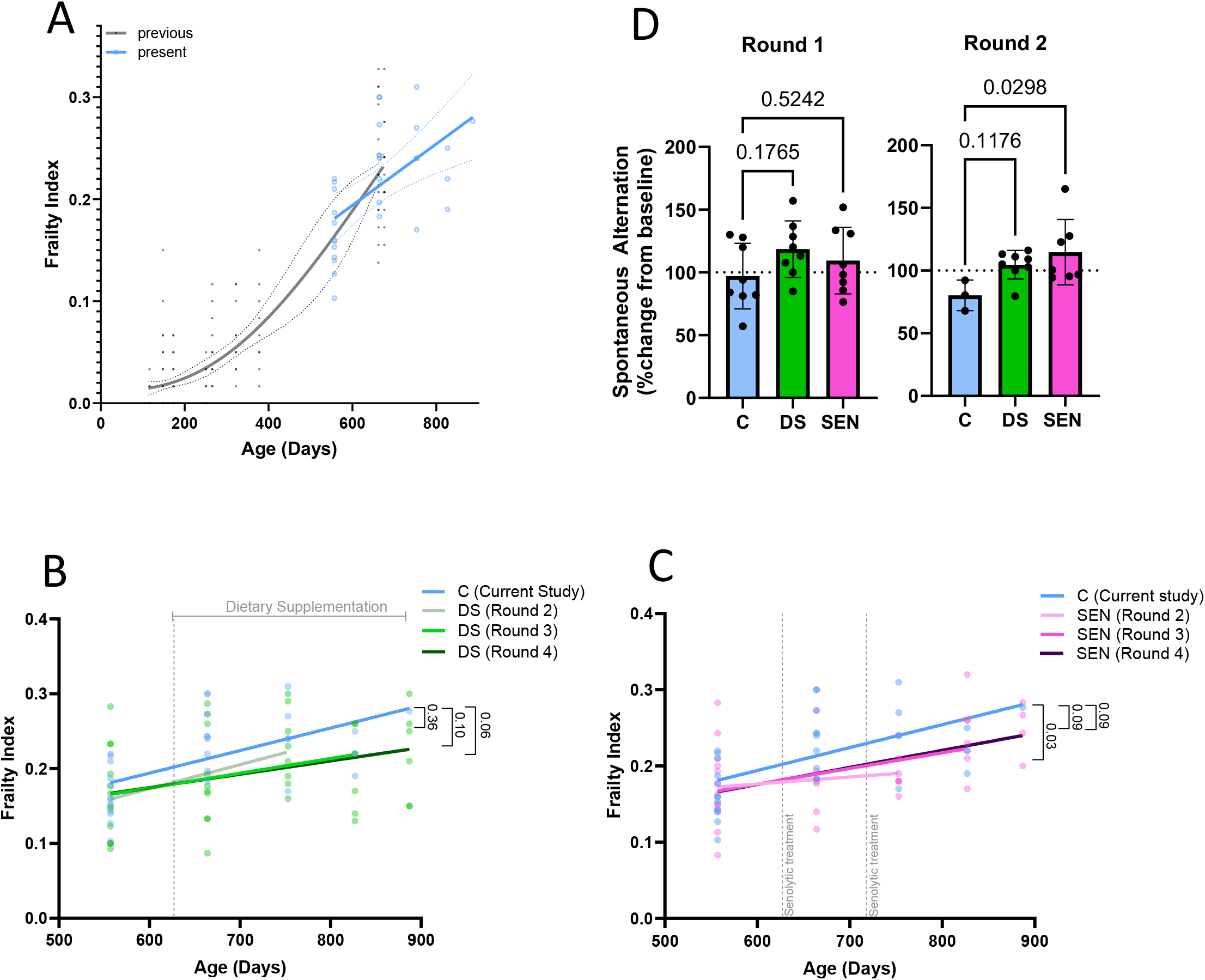
Health span effects. **A)** Comparison between frailty indices in present (blue) and previous [28] (black) controls. All animals were male C57Bl/6J mice. Individual frailty indices (dots), regression lines (solid lines) and 95% confidence intervals (dotted lines) are shown. Frailty indices in the DS **(B)** and SEN **(C)** group in comparison to controls. Frailty indices for each animal (dots) and linear regression lines from baseline to every assessment time point are indicated to illustrate the kinetic developments. p values for the differences in regression slopes to that of control are shown. **D)** Spontaneous alternation in a Y-maze 1 month following the first and second round of senolytic treatment as percentage change from each mouse’s pre-treatment performance. Percentage changes of individual mice are illustrated as dots. Data are mean ± SD, n≥3. ANOVA p values for differences between controls and treatment groups are indicated.

### 4. The multi-component NOVOS supplement has an effect on senescent cell size, but not senolytic activity in human fibroblasts

We tested the senolytic activity of the NOVOS multi-ingredient supplement as a whole and its individual ingredients in human fibroblasts to reveal potential mechanisms of its anti-ageing activity. This was with the exception of Calcium alpha-ketoglutarate, which did not adequately solubilise. Over a wide range of concentrations, neither the multi-component supplement (Fig. 3a) nor any of the tested ingredients (Figs. 3b-j) showed a preferential reduction of senescent over non-senescent fibroblast viability. This suggests a lack of senolytic effect from the supplement, despite the inclusion of Fisetin, which has previously been reported as a (weak) senolytic in mouse ERCC1^-/-^ fibroblasts and IMR90 human fibroblasts^6^.

**Fig. 3.**
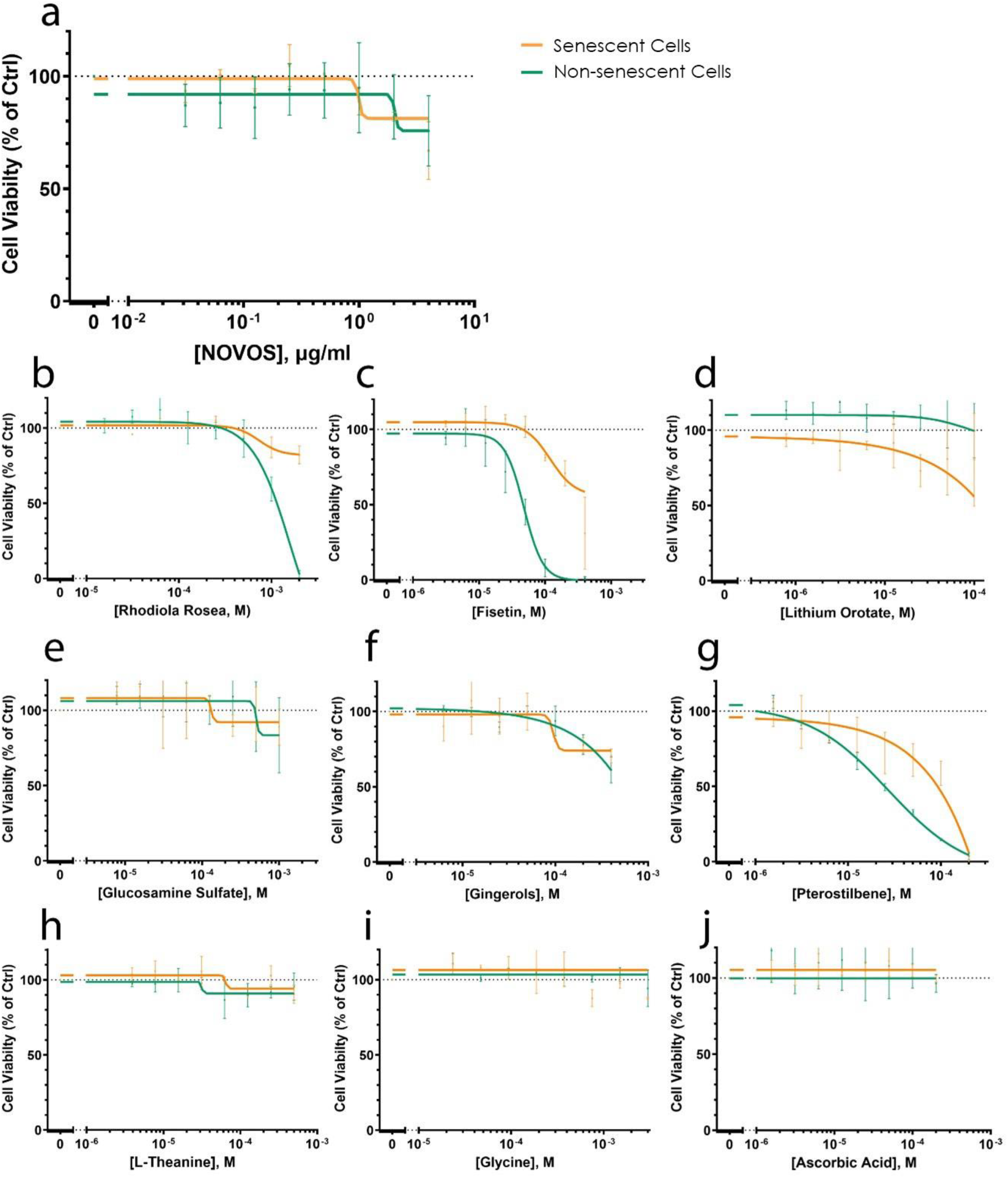
The multi-ingredient NOVOS supplement or its components do not act as senolytic in human fibroblasts. Senescent (orange) and non-senescent (green) human fibroblasts were treated for 3 days with the indicated concentrations of NOVOS supplement **(a)** or its active ingredients **(b-j)** and cell viability was measured using crystal violet. Titration curves are presented as moving averages ± SEM from between 2 and 4 experiments per compound.

Using co-cultured senescent and non-senescent cells, we tested serial dilutions of the multi-ingredient supplement on senolytic capacity and cell size (Figs. 4a-b). Cell size can be indicative of a senostatic activity, as illustrated with the senostatic effects of rapamycin (Supplementary Fig. S1). Because of the diverse solubilities of the ingredients, we prepared stock solutions of the multi-ingredient supplement in both water and ethanol and tested these separately. Confirming the previous results, we did not see preferential ablation of senescent cells by the complete supplement (Figs. 4a-d), whether it was dissolved in ethanol (Fig. 4c) or water (Fig. 4d). However, size of senescent cells was decreased preferentially by the complete supplement when dissolved in ethanol (Figs. 4b,e) compared to the water dissolved complete supplement (Figs. 4d,f). Given that Fisetin on its own is much less soluble in water than in ethanol, and its previously reported effects on senescence we investigated this alone in the co-culture assay. Again, Fisetin alone again did not show a senolytic effect, however, there was an effect on cell size in senescent cells between 1µM and 10µM (Fig. 4g).

**Fig. 4.**
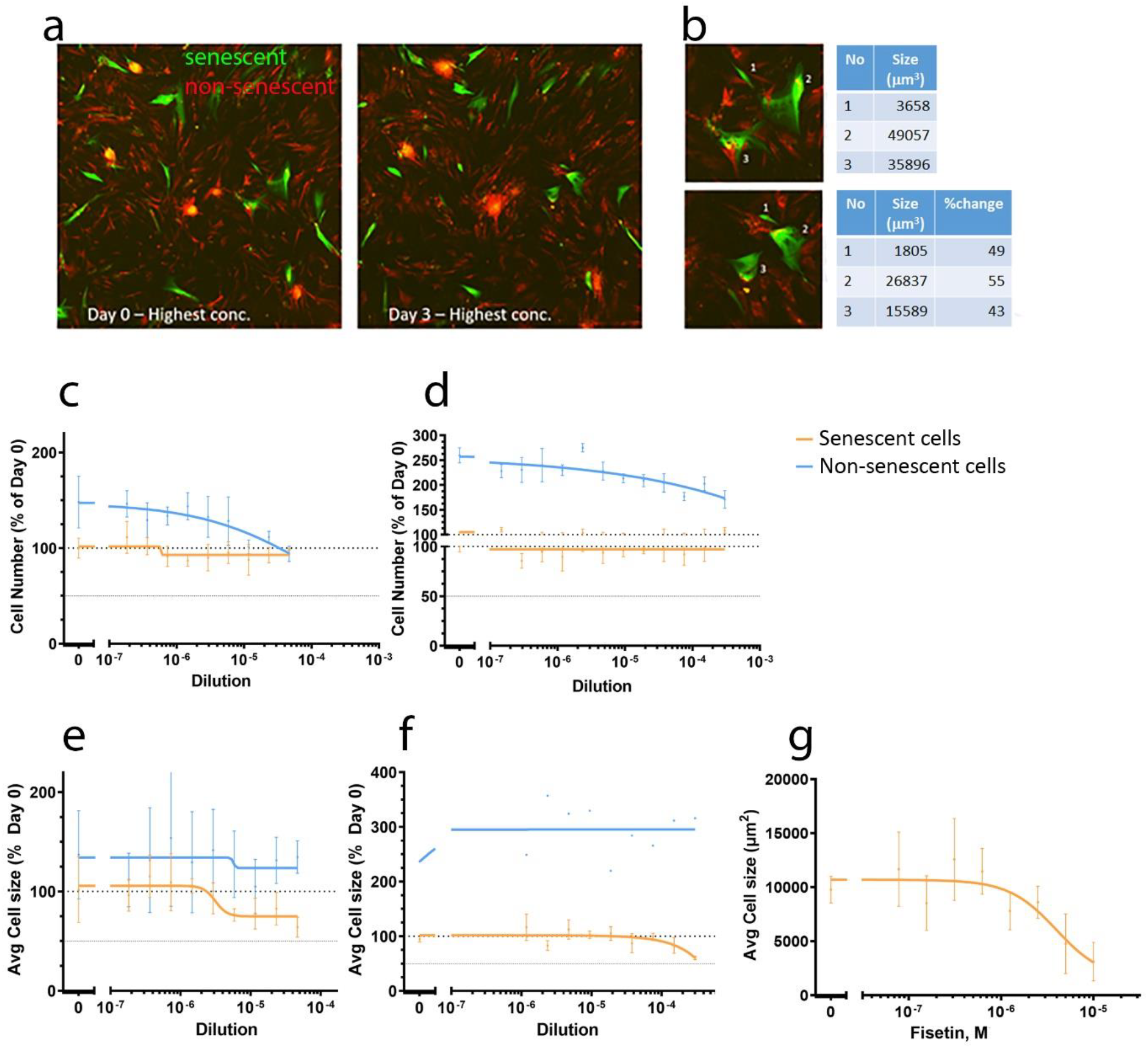
The multi-ingredient NOVOS supplement and its component fisetin act as senostatic in human fibroblasts. **a)** Representative micrographs of the senescent/non-senescent human fibroblast co-culture before (day 0) and after (day 3) treatment with the complete NOVOS supplement at 5×10^−5^ dilution. Senescent cells are labelled in green and non-senescent in red. **b)** Higher magnification images of three senescent cells from **(a)** at start (top) and end (bottom) of treatment with their respective areas indicated. **c)** Effect of the complete NOVOS supplement diluted in ethanol on numbers of senescent (orange) and non-senescent (blue) cells. **d)** Effect of the complete NOVOS supplement diluted in water on numbers of senescent (orange) and non-senescent (blue) cells. **e)** Effect of the complete NOVOS supplement diluted in ethanol on average sizes of senescent (orange) and non-senescent (blue) cells. **f)** Effect of the complete NOVOS supplement diluted in water on average size of senescent (orange) and non-senescent (blue) cells. **g)** Effect of Fisetin diluted in DMSO on average size of senescent cells. Data are mean ± SEM from 3 technical repeats.

## Discussion

Both the multi-ingredient dietary supplement and the senolytic intervention tended to increase lifespan of male mice (median lifespan increased by 18 and 21% over controls, respectively). However, control lifespans in the present study (703 days) were shorter than in our previous study on the same mouse strain (median lifespan of 810 days)^27^. Two factors may account for this difference. First, mice in the present study received soaked food from 18 months of age, resulting in higher food intake, which caused on average 5 to 10g higher body weights compared to mice fed standard food pellets at the same age. There is a fine balance between health-deteriorating consequences of being overweight and health-promoting effects of body weight maintenance in older mice, and it is possible that this effect contributed to the shorter lifespan of controls. Additionally, there was a technical confounding factor at the first round of treatment; using intraperitoneal injections as the route of administration of BAM15 (or vehicle), which likely resulted in a wave of deaths across all three groups within the two weeks following treatment. The postmortem examinations suggested that these deaths were likely due to infection, namely the seminal vesicles which become enlarged with age were nicked with the needle upon entry. If those deaths were censored, the resulting median lifespan in the controls equalled 719 days, the DS group equalled 860.5 days, and SEN group equalled 852 days (Supplementary Fig. S4). Following censoring, the median lifespans of the DS and the SEN group were 19.7% and 18.5% longer than the controls respectively. This suggests that in relative terms our results are robust to this confounding factor.

Effects of the interventions on healthspan measures were dependent on the treatment schedules. In the DS group, the progression of frailty with age was slowed down as the intervention continued and differences became almost significant after 10 months of treatment over controls. Short-term memory, as assessed by spontaneous alternation, in the mice assigned to the DS intervention was sustained, whereas the Controls declined over time. Conversely, senolytic effects on frailty and cognition were significant shortly after the second round of treatment but waned over the next three months. This is in some contrast to previous results, where either first-generation senolytics^29^ or the same senolytic treatment using Navitoclax and BAM15 as in the present study^23^ were given to mice after sublethal irradiation. Irradiation can cause a rapid increase in senescence burden, resulting in around a two-fold enhanced progression rate of frailty^24^. In these cases, a single round of senolytics was sufficient to normalize frailty progression to control levels for at least one year^23,29^. However, the irradiated mice were not studied beyond 17 months of age, and it is probable that the continuous generation and accumulation of senescent cells at higher ages requires repeated senolytic interventions. This could be easily done in future because the dose of the potentially toxic senolytic Navitoclax has been reduced by two orders of magnitude by combination with the mitochondrial uncoupler BAM15^23^.

There was no evidence of senolytic activity of the NOVOS multi-ingredient supplement or its individual ingredients in human fibroblasts. Given the heterogeneity of the senescent phenotype^30^, our results cannot exclude that the NOVOS multi-ingredient supplement or some of its ingredients may have senolytic activity towards other cell types. However, our screens suggested a senostatic activity for the NOVOS multi-ingredient supplement and its ingredient, Fisetin.

Treatment of old mice with 500mg/kg BW Fisetin have previously shown an increase in lifespan of approximately 10%^6^, with short-term treatment reducing senescent cell numbers at 100mg/kg and improved their arterial function^31^. In our intervention with the NOVOS multi-ingredient supplement, Fisetin would have been given to the mice at 18mg/kg BW, a much lower dose, and likely does not account fully for the lifespan extension seen. However, comparable oral route doses (20mg/kg) in mice have previously shown some positive effects on object recognition memory^32^ and in protecting against virus induced mortality^33^. Previously we had shown that the net effect of a continuous senostatic intervention, dietary restriction, in immunocompetent mice was a reduction of senescent cell frequencies because it inhibited de-novo generation of senescent cells from pre-existent ones via bystander effects^34^. Therefore, senostatic effects might be one mechanism contributing towards the positive life- and health-span effects of the NOVOS dietary supplement.

The median lifespan extension we saw here was larger than that previously seen with Fisetin, while this may be from the combination of ingredients, our findings are complicated by the loss of mice to presumed infection following IP injection during the first intervention point. Our study was also limited solely to male mice due to funding.

However, to our knowledge it is the first study to directly compare health- and life-span effects of a defined multi-ingredient nutraceutical with a senolytic treatment. Both interventions, despite being started only at relatively advanced age, in ‘overweight’ animals, improved survival over controls and showed promises with respect to frailty and cognitive health. Our study provides clear indications for the requirements of optimisation of treatment regimens for each specific intervention and a strong rationale for further work to explore the mechanisms of action of multi-ingredient nutraceuticals.

