## Supplementary Figs. and Tables for "Comparable anti-ageing efficacies of a multi-ingredient nutraceutical and a senolytic intervention in old mice"

**Supplementary Figures**

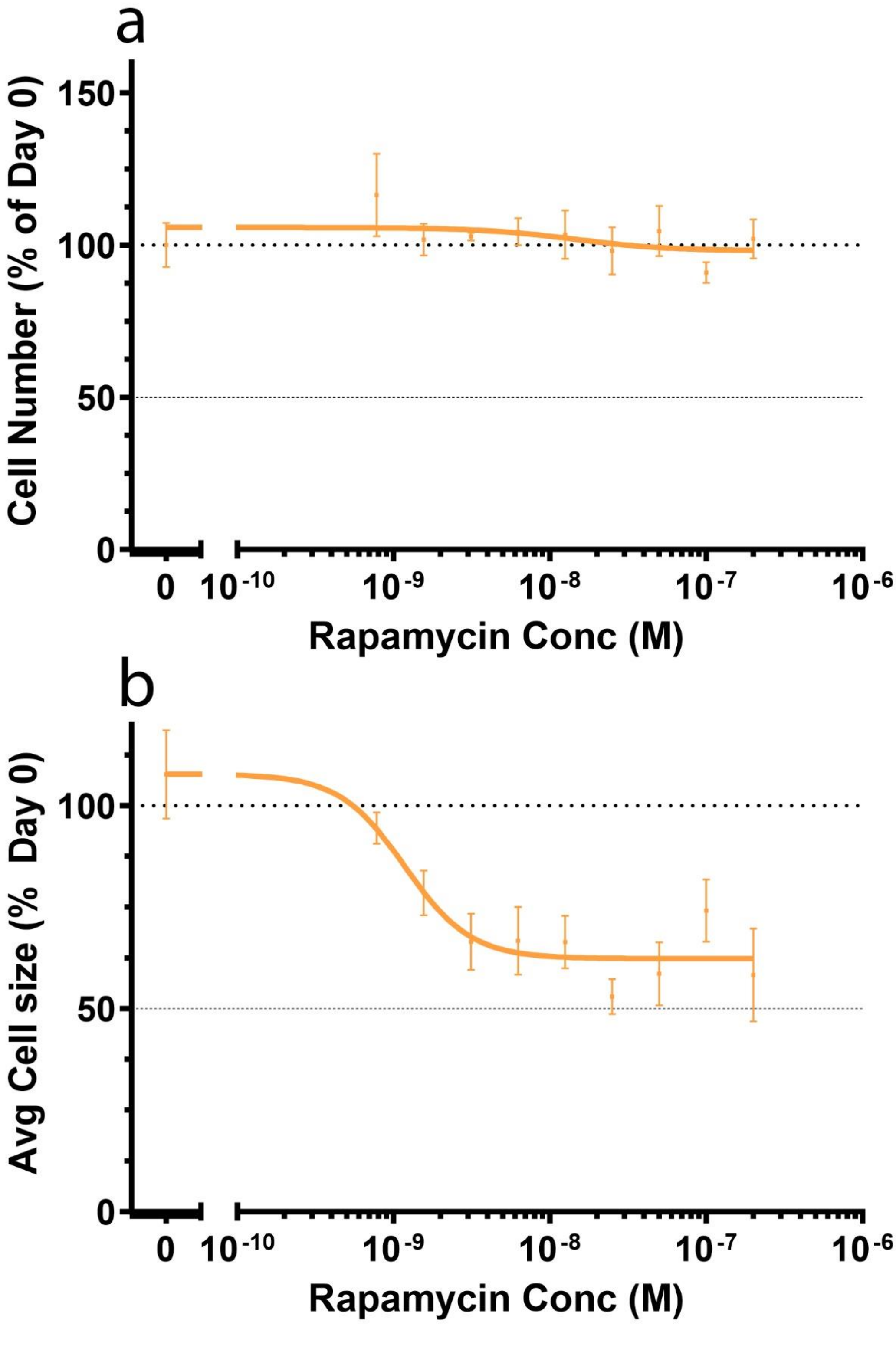


**Fig. S1: Effect of the senostatic rapamycin on number (A) and size (B) of human fibroblasts.** Data are mean ± SEM from 3 technical repeats.

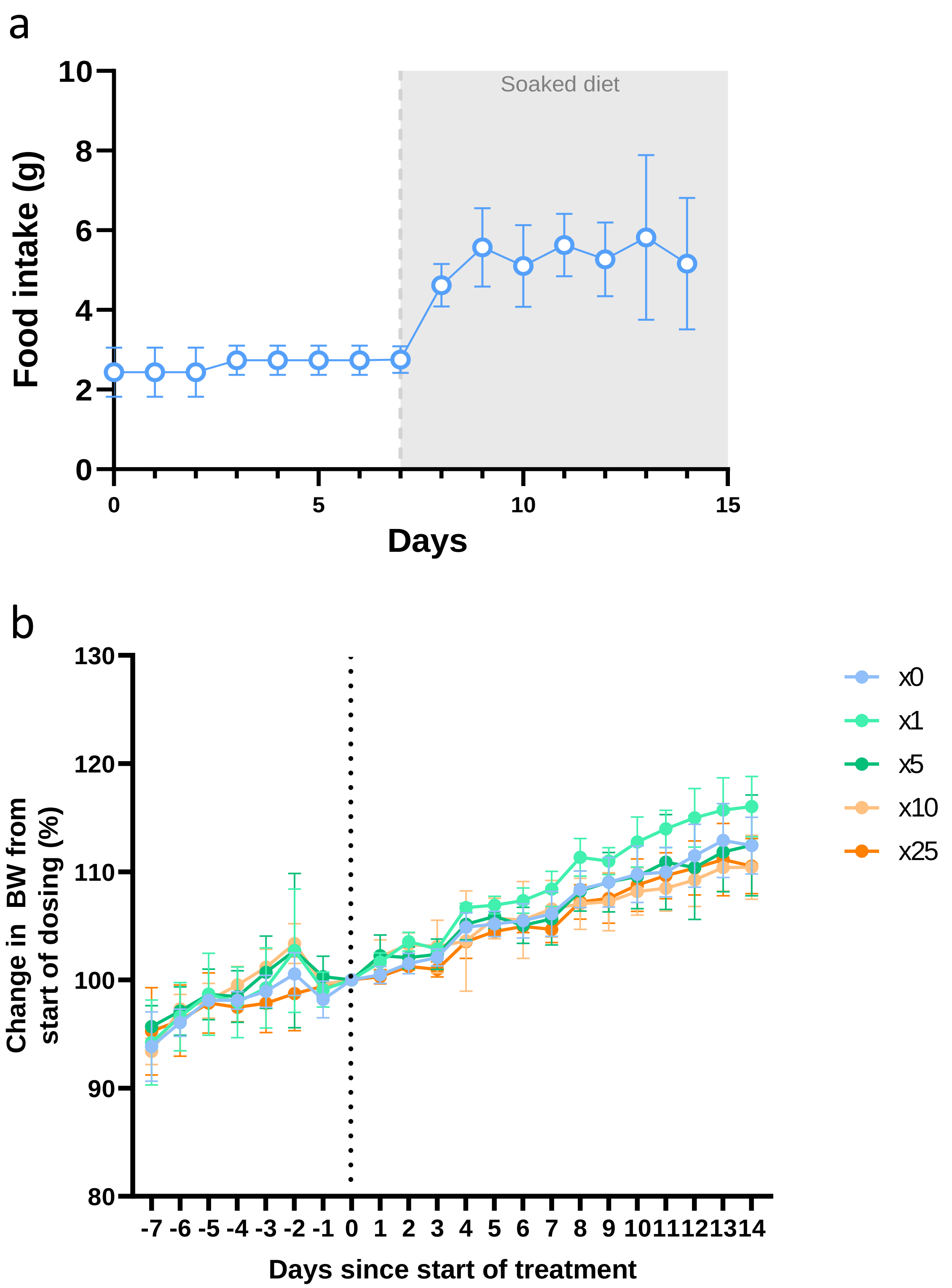


**Fig. S2: Food intake and BW in the dose finding study. A)** Food intake before and after a switch from dry pellets to soaked food. Data are mean± SD, n=20. **B)** Mice were switched at day -7 to soaked food, which was supplemented with the multi-component nutraceutical in the indicated doses from day 0. Data are mean ± SD, n=4/group.

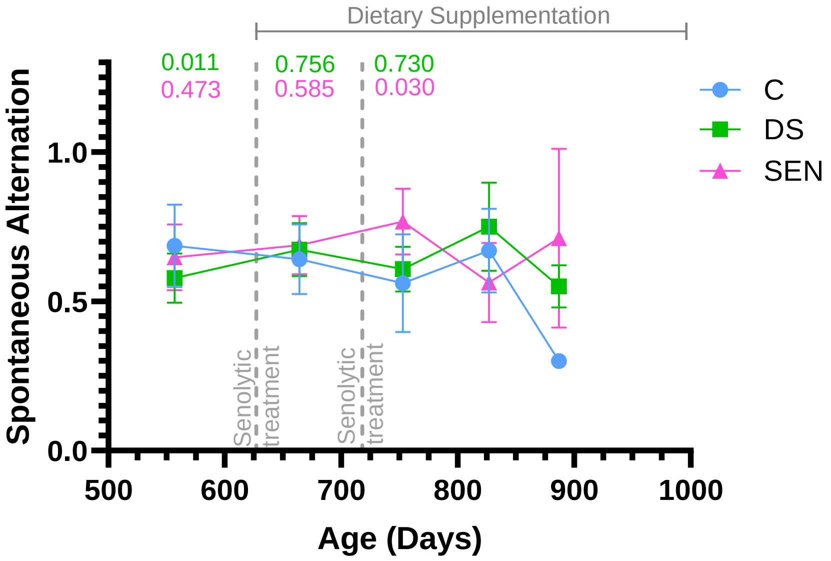


**Fig. S3: Changes of short-term memory.** Short-term memory assessed by Y-maze spontaneous alternation. Controls (blue), DS (green), SEN (pink). Data are mean ± SD, n≥3, except for Round 3 and Round 4 which contained 2 and 1 Control respectively. ANOVA p values for differences between controls and colour-coded treatment group are indicated for Baseline, Round 1, and Round 2 assessment time points.

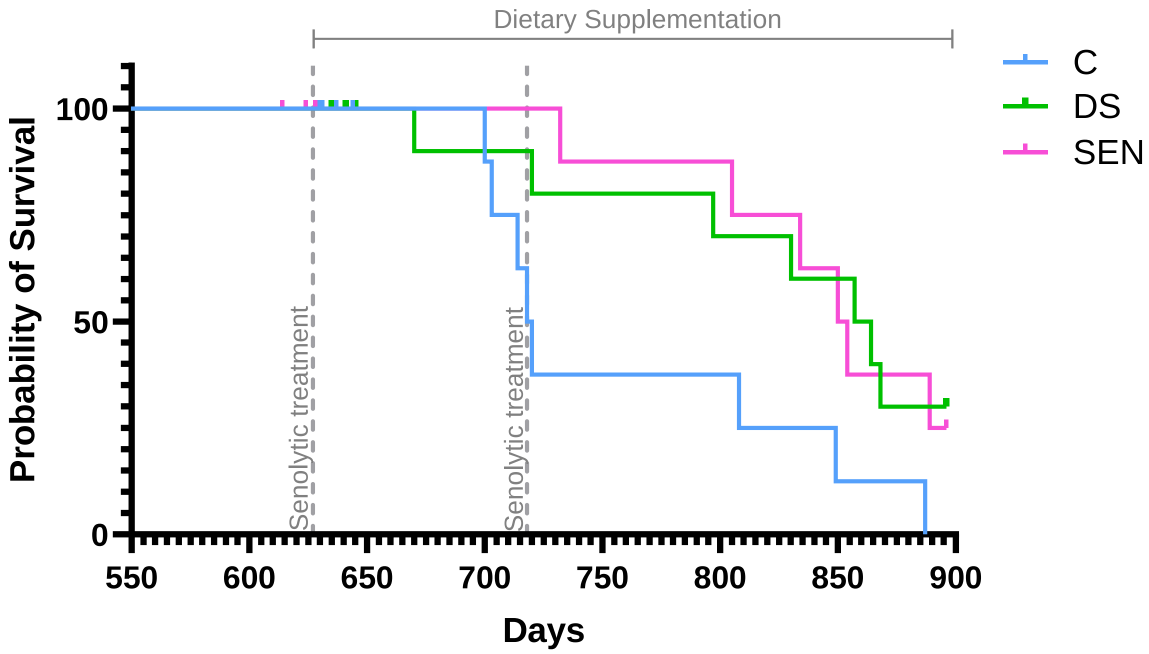


**Fig. S4: Survival after censoring for infection-induced deaths.**

Kaplan-Meier survival curves. Median survival for C = 719 (700,850), DS = 860.5 (670, 894), and S = 852 (730, 900). Log-rank tests for the difference to controls resulted in p=0.044 (DS) and p=0.020 (SEN), respectively.

**Supplementary Tables**

**Table S1: Death record of mice throughout study.**

| Mouse | Study Group | Birth Date | Date of Death | Age at Death (months) | Cause of Death |
|---|---|---|---|---|---|
| 15 LN | Senolytic | 04/11/2021 | 11/07/2023 | 20 | Seizure |
| 20 LN | Senolytic | 04/11/2021 | 21/07/2023 | 20 | Distended abdomen |
| 13 NN | Senolytic | 04/11/2021 | 25/07/2023 | 20 | Moribund |
| 27 LN | Control | 04/11/2021 | 27/07/2023 | 20 | Enlarged Seminal Vesicle |
| 18 LN | Control | 04/11/2021 | 28/07/2023 | 20 | Enlarged Seminal Vesicle |
| 17 LN | NOVOS | 04/11/2021 | 01/08/2023 | 21 | Enlarged Seminal Vesicle |
| 11 RN | Control | 04/11/2021 | 03/08/2023 | 21 | Enlarged liver and blocked bladder |
| 14 NN | Senolytic | 04/11/2021 | 03/08/2023 | 21 | Continued weight loss |
| 22 RN | NOVOS | 04/11/2021 | 07/08/2023 | 21 | Seminal Vesicle tumour |
| 282 NN | Control | 17/11/2021 | 10/08/2023 | 21 | Enlarged seminal vesicle |
| 26 LN | NOVOS | 04/11/2021 | 11/08/2023 | 21 | Seminal Vesicle tumour |
| 24 LN | Senolytic | 04/11/2021 | 11/08/2023 | 21 | Pancreatic tumour |
| 12 NN | NOVOS | 04/11/2021 | 05/09/2023 | 22 | Severe dermatitis |
| 27 NN | Control | 04/11/2021 | 05/10/2023 | 23 | Intranasal mass |
| 18 NN | Control | 04/11/2021 | 08/10/2023 | 23 | Continued weight loss |
| 21 NN | Control | 04/11/2021 | 19/10/2023 | 23 | Incisor mass |
| 21 RN | Control | 04/11/2021 | 23/10/2023 | 23 | Liver tumour |
| 282 RN | Control | 17/11/2021 | 25/10/2023 | 23 | Swelling and discharge from eye |
| 17 RN | NOVOS | 04/11/2021 | 25/10/2023 | 23 | Moribund |
| 24 NN | Senolytic | 04/11/2021 | 06/11/2023 | 24 | Enlarged seminal vesicle |
| 26 RN | NOVOS | 04/11/2021 | 10/01/2024 | 26 | Possible stroke |
| 16 RN | Senolytic | 04/11/2021 | 18/01/2024 | 26 | Seizure |
| 27 RN | Control | 04/11/2021 | 21/01/2024 | 26 | Lung tumour |
| 17 NN | NOVOS | 04/11/2021 | 12/02/2024 | 27 | Continued weight loss |
| 16 NN | Senolytic | 04/11/2021 | 16/02/2024 | 27 | Ulcerated tail wound |
| 11 LN | Control | 04/11/2021 | 02/03/2024 | 28 | Overall frailty |
| 19 RN | Senolytic | 04/11/2021 | 03/03/2024 | 28 | Continued weight loss |
| 20 RN | Senolytic | 04/11/2021 | 07/03/2024 | 28 | Large intestine tumour |
| 70 RN | NOVOS | 17/11/2021 | 10/03/2024 | 28 | Continued weight loss |
| 22 NN | NOVOS | 04/11/2021 | 17/03/2024 | 28 | Moribund |
| 22 LN | NOVOS | 04/11/2021 | 21/03/2024 | 28 | Continued weight loss |
| 282 LN | Control | 17/11/2021 | 09/04/2024 | 29 | Seizure |
| 16 LN | Senolytic | 04/11/2021 | 11/04/2024 | 29 | Continued weight loss |

**Table S2: Composition of the NOVOS multi-ingredient dietary supplement per recommended daily dose for humans** (assuming 70kg body weight)

| Ingredients | |
|---|---|
| **Vitamin C** | 100mg |
| **Mag Malate** | 304mg |
| **Magnesium** | 231mg |
| **Malate** | 2000mg |
| **Glycine** | 1700mg |
| **Calcium Alpha Ketglutaric Acid** | 1100mg |
| **Glucosamine Sulphate** | 1000mg |
| **Rhodiola Rosea Root extract** | 300mg |
| **L-Theanine** | 150mg |
| **Hyaluronic acid** | 100mg |
| **Fisetin** | 100mg |
| **Ginger Root Extract** | 100mg |
| **Pterostilbene** | 50mg |
| **Lithium Orotate** | 20mg |

**Table S3. Stock composition and maximum tested concentrations for the senolytic assays**

| Compound | Stock concentration | solvent | Maximum tested concentration |
|---|---|---|---|
| NOVOS | 0.5 mg/ml | DMSO: H2O | 4ug/ml |
| *Rhodiola Rosea* | 500 mM | H2O | 2mM |
| Fisetin | 100 mM | DMSO | 0.4 mM |
| Lithium Orotate | 25 mM | DMSO | 0.1 mM |
| Glucosamine Sulfate | 500 mM | DMSO | 1 mM |
| Gingerols | 100 mM | DMSO | 0.4 mM |
| Pterostilbene | 50 mM | EtOH | 0.2 mM |
| L-Theanine | 250 mM | H2O | 0.5 mM |
| Glycine | 30 mM | H2O | 3 mM |
| Ascorbic Acid | 100 mM | H2O | 0.2 mM |

**Table S4: Composition of the NOVOS multi-ingredient dietary supplement per 12.3x dose of 40g mouse**

| Ingredients | Intake of a mouse at 40g weight (mg/kg) |
|---|---|
| **Vitamin C** | 18.0 |
| **Mag Malate** | 361.0 |
| **Magnesium** | 54.1 |
| **Malate** | 306.8 |
| **Glycine** | 361.0 |
| **Calcium Alpha Ketglutaric Acid** | 198.5 |
| **Glucosamine Sulphate** | 180.5 |
| **Rhodiola Rosea Root extract** | 54.1 |
| **L-Theanine** | 27.1 |
| **Hyaluronic acid** | 18.0 |
| **Fisetin** | 18.0 |
| **Ginger Root Extract** | 14.4 |
| **Pterostilbene** | 9.0 |
| **Lithium Orotate** | 4.7 |
